# The urinary microbiota composition remains stable over time and under various storage conditions

**DOI:** 10.1101/2019.12.13.875187

**Authors:** Caspar Bundgaard-Nielsen, Nadia Ammitzbøll, Yusuf Abdi Isse, Abdisalam Muqtar, Ann-Maria Jensen, Peter Leutscher, Louise Thomsen Schmidt Arenholt, Søren Hagstrøm, Suzette Sørensen

## Abstract

**Background:** New sensitive techniques have revealed a large population of bacteria in the human urinary tract, challenging the perception of the urine of healthy humans being sterile. While the role of this urinary microbiota is unknown, dysbiosis has been linked to disorders like urgency urinary incontinence and interstitial cystitis. When comparing studies it is crucial to account for possible confounders introduced due to methodological differences. Here we investigated whether storage condition or time of collection, had any impact on the urinary microbial composition.

**Results:** For comparison of different storage conditions, urine was collected from five healthy adult female donors, and analyzed by 16S rRNA gene sequencing. Using the same methods, the daily or day-to-day variation in urinary microbiota was investigated in nineteen healthy donors, including four women, five men, five girls, and five boys. With the exception of two male adult donors, none of the tested conditions gave rise to significant differences in alpha and beta diversities between individuals. *Conclusion*: The composition of the urinary microbiota was found to be highly resilient to changes introduced by storage temperature and duration. In addition, we did not observe any intrapersonal daily or day-to-day variations in microbiota composition in women, girls or boys.

Together our study supports flexibility in study design, when conducting urinary microbiota studies.

**Author summary:** The discovery of bacteria native to the urinary tract in healthy people, a location previously believed to be sterile, has prompted research into the clinical potential of these bacteria. However, methodological weaknesses can significantly influence such studies, and thus development of robust techniques for investigating these bacteria are needed. In the present study, we investigated whether differences in storage following collection, could affect the bacterial composition of urine samples. Next, we investigated if this composition exhibited daily or day-to-day variations.

Firstly, we found, that the bacterial composition of urine could be maintained by storage at −80 °C, −20 °C, or refrigerated at 4 °C. Secondly, the bacterial composition of urine remained stable over time. Overall, the results of this study provide information important to study design in future investigations into the clinical implications of urinary bacteria.

## Background

It has been established that the human body has a symbiotic relationship with an abundance of microorganisms, which play a role in maintenance of health. In particular, microorganisms present in the gut have received much attention, and many studies have described their beneficial functions in immune regulation[1,2] and metabolic processes[3]. However, when brought out of balance (dysbiosis), the same microorganisms have been associated with several pathological states including infections[4], autoimmune diseases[5], obesity[6], and psychiatric or neurodevelopmental disorders[7–9].

Until recently, it was believed that urine under normal conditions was sterile. This has now been challenged by sensitive PCR-based techniques including 16S rRNA gene sequencing and expanded quantitative urine culture (EQUC). Studies are now emerging, investigating the urinary microbiota in various patient groups and healthy participants, showing that urine contains a plethora of bacteria with yet unknown function[10–12]. Most studies published on urinary microbiota are primarily on women, fewer on men[13] and only a single on young children[14]. In general, the core microbiota composition is very different between women and men. The urine of women is mainly dominated by *Lactobacillus* followed by *Gardnerella* genera[11,13,15–17]. Men have a more diverse bacterial composition consisting of broader representation of different genera including *Lactobacillus*, *Corynebacterium*, *Staphylococcus*, and *Prevotella*[13,18–23]. It is assumed that these bacteria provide healthy functioning of the lower urinary tract. Notably, several studies have recently documented that an alteration in urinary microbiota correlates to diseases in the lower urinary tract, including urgency urinary incontinence[10–12,24] and interstitial cystitis[15,25]. More knowledge on the urinary microbiota may therefore help us to understand the etiology behind diseases of the lower urinary tract.

Despite the growing interest for urinary microbiota research, it appears that the methodologies and study designs, used in different studies, are highly heterogeneous, which makes it difficult to interpret and compare observed findings. Several protocol optimization studies have been conducted on fecal samples, providing valuable guidelines on how to store and process samples for gut microbiota studies[26–31]. Importantly, the chemical content and structure of urine is very different from feces, leaving urinary microbiota research as a bare and unexplored field regarding protocol recommendations. In fact, few studies have investigated the technical and methodological aspects of urine microbiota research[19,32–34]. These mainly focused on the urine collection method. e.g., suprapubic aspiration, clean-catch midstream, or transurethral catheterization sample collection. However, only a single study investigated how different temperatures and use of a stabilization buffer could affect the urinary microbiota in healthy women[34]. None of the studies has taken into consideration whether the microbiota remains stable over time.

We aimed to determine if different urine storage temperatures could influence microbiota composition in healthy women. Furthermore, we investigated if the urinary microbiota remained stable throughout the day or between two different days in healthy women, men, and children.

## Results

### Different storage conditions do not critically affect bacterial composition

Due to the risk of DNA degradation or bacterial growth, the ideal sampling strategy for urine microbiota analyses would be to purify DNA immediately following urination, or to transfer the urine samples directly to −80 °C or colder. This is however not always possible or practical in a clinical setting, or when utilizing self-sampling at the home of the study participants. We therefore tested if storage of urine at different sub-optimal temperatures, altered the microbiota composition compared to a freshly processed sample. For this purpose, urine was collected from five healthy donors. Each urine sample was subsequently divided and stored according to one of the seven combinations of temperatures and times (Fig 1). Since each condition was tested in duplicate experiments, we reached a total of 70 samples. After the allocated storage period, total DNA was purified, and DNA concentrations obtained ranged from 16.2-248.0 ng/mL urine with minor variations between donors (Fig 2). With the exception of donor E, the highest DNA concentration was observed in the freshly processed sample, followed by samples stored directly at −80 °C.

**Fig 1:**
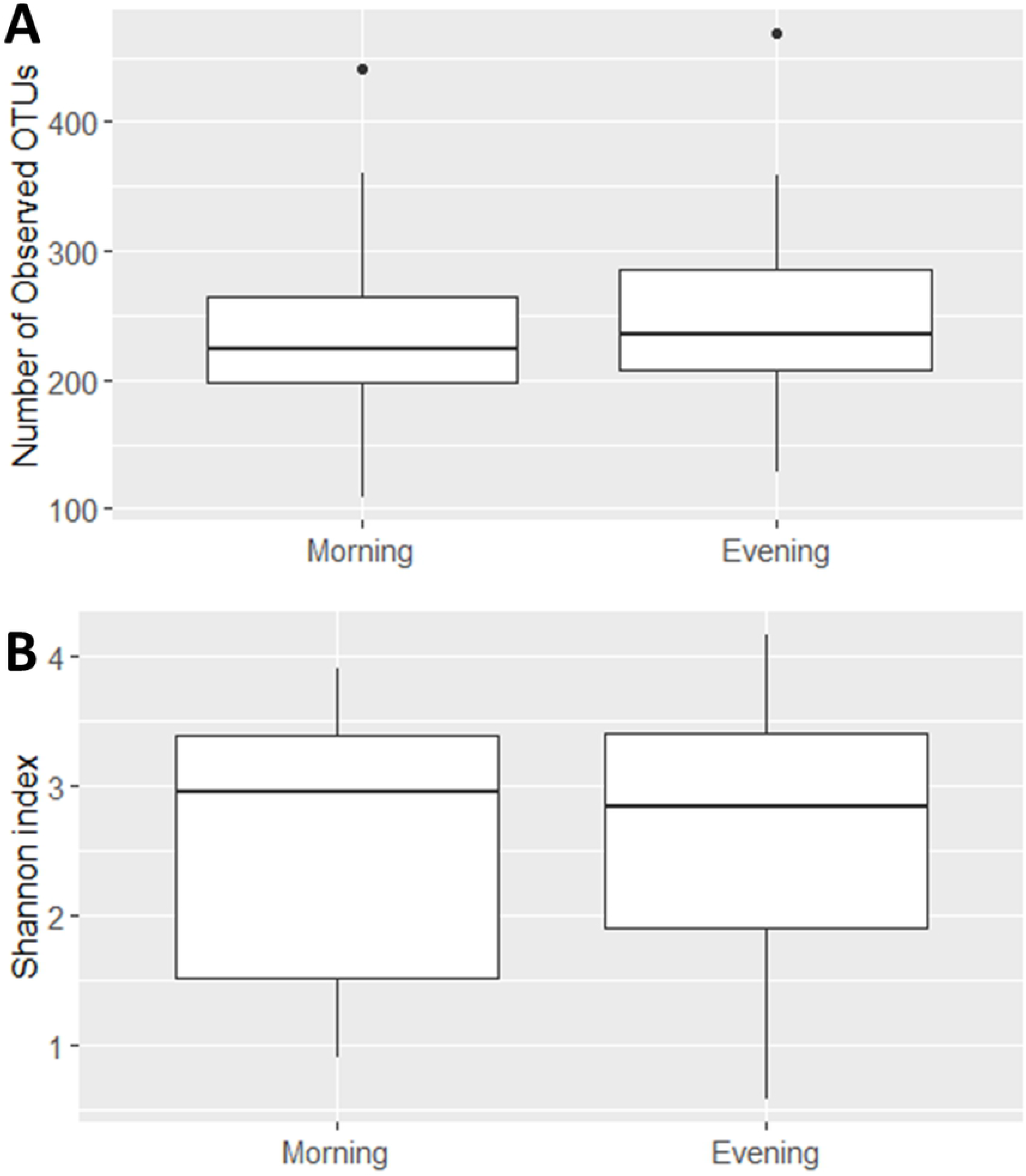
Urine storage conditions. Urine samples were either processed directly (room temperature, 0 hours) or at different temperatures for different time periods. Duplicate experiments were performed.

**Fig 2:**
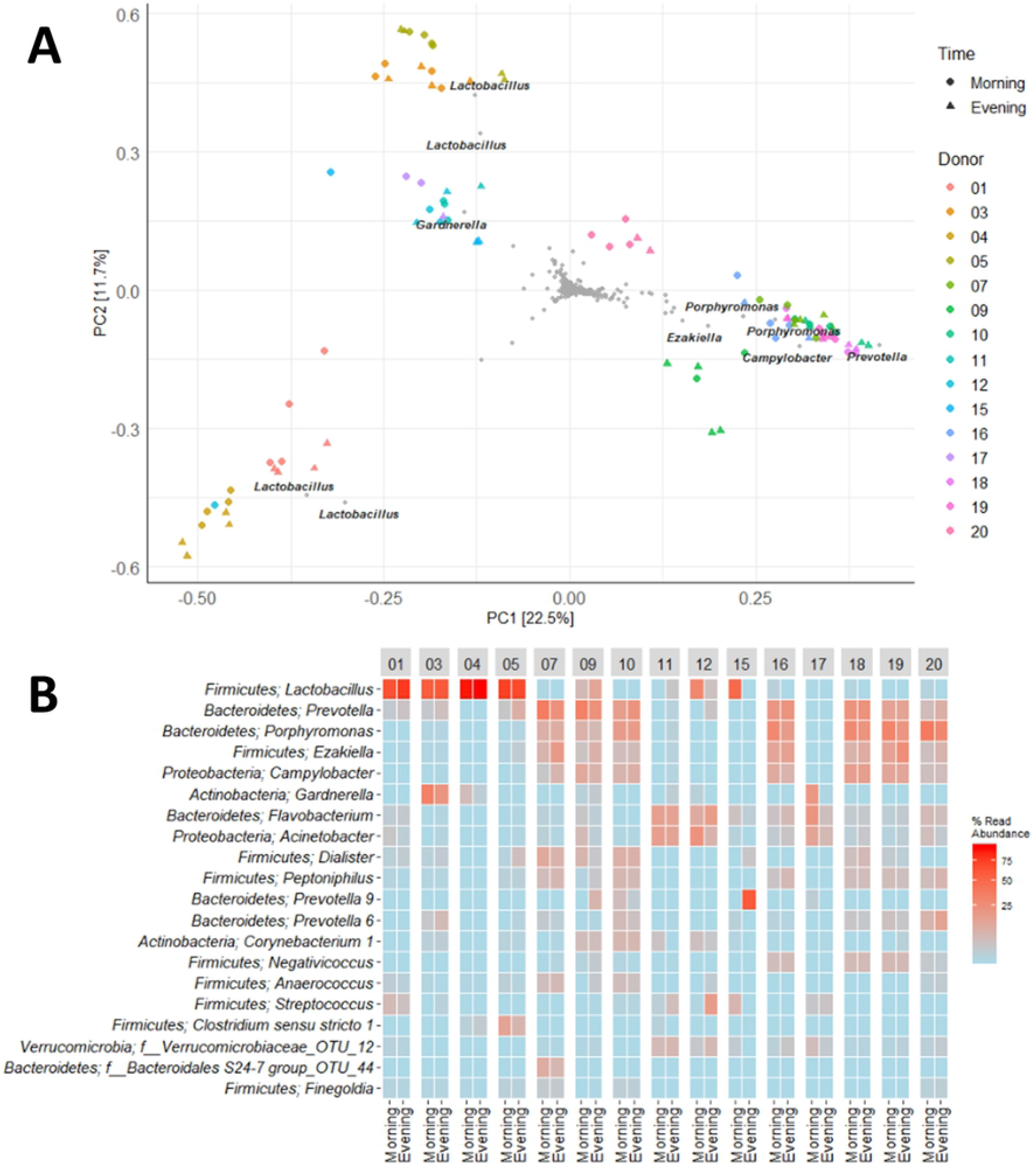
DNA yield. Quantity of total DNA purified from urine shown for each donor (A-E) and storage condition. Each sample is named as “temperature”_”duration”. *P<0.05. RT = room temperature, >72 = sample are stored for at least 72 hours.

16S rRNA gene sequencing of the V4 region resulted in a total of 2,097,325 reads (median 29,334, range 1299-132,881 reads per sample). 933 unique Operational taxonomic units (OTUs) were identified (median 106, range 33 - 502 OTUs per sample), with taxonomy assigned on the phylum level for 96.1 % of OTUs and genus level for 53.2 %. A rarefaction curve was generated, and used to deselect samples that did not adequately cover all unique OTUs and therefore showed insufficient sequencing coverage (Fig 3A). Consequently, three samples were removed. Four samples were furthermore discarded based on poor duplicate comparison (Fig 3B and 3C), probably caused by background contamination due to low-biomass samples. This led to a total of 63 samples being included in the following analyses.

**Fig 3:**
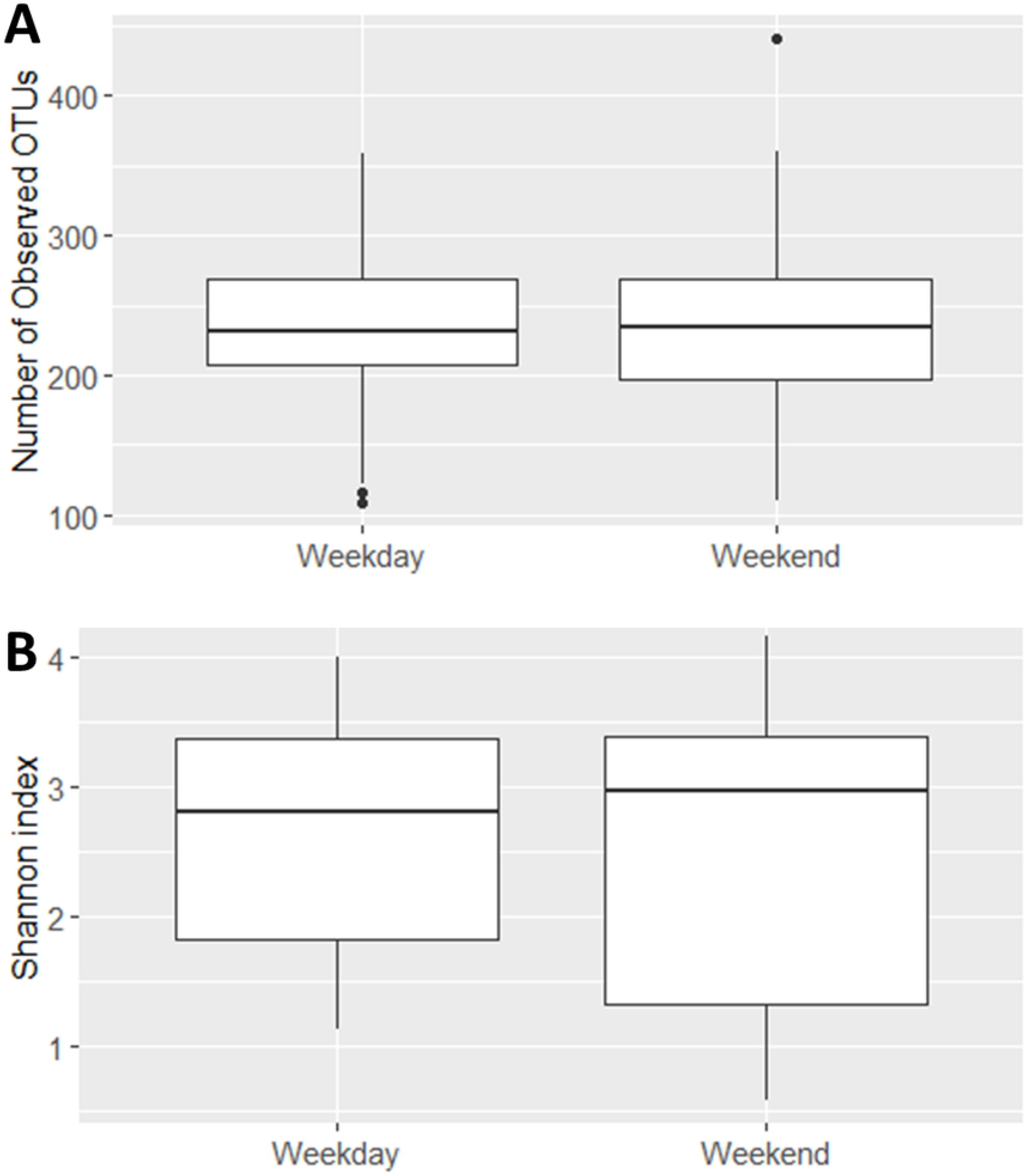
Quality control used for removing low quality samples. A. Rarefaction Curve, showing number of unique OTUs generated based on quantity of reads. The red dotted line indicates cutoff value of 10.000 reads. Three samples were excluded due to values below cutoff. B. Principal Component Analysis (PCA) clustering of samples using Hellinger transformed OTU abundances, in order to identify possible outliers. C. Heatmap depicting the 20 most common OTUs for each separate duplicate. Each name consists of phylum followed by genus name. If no genus could be identified, the best taxonomic assignment is listed. The heatmap was used to exclude potential outliers. The following four samples were excluded: A −20 72h_2, D −20 72h_2, D 4 24h_1 and E −80 >72h_2. These has been circled in B. and marked with a red * in C.

Different storage conditions did not result in significant differences in alpha diversity metrics, including OTU richness (Fig 4A) and Shannon diversity (Fig 4B). This indicated that bacterial growth was limited. Interestingly, when looking at beta diversity it appears that variations between storage conditions are minor compared to interpersonal variations (Fig 5). This supports, therefore the validity of using other storage conditions, for urine microbiota analyses, than normal gold standard conditions.

**Fig 4:**
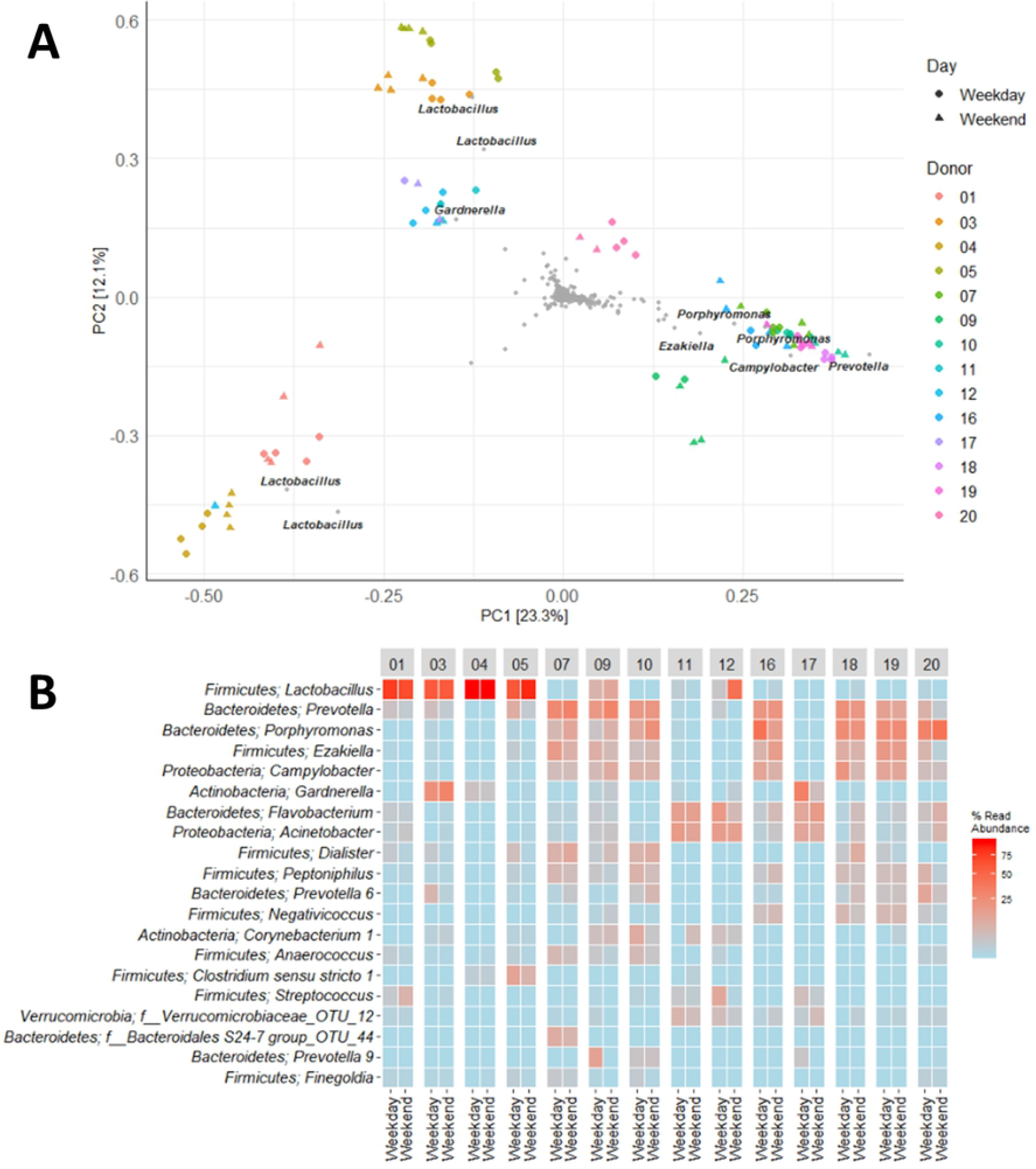
Effects of storage temperature and duration on urinary microbiota alpha diversity. A. OTU richness showing numbers of unique OTUs observed in different storage conditions and durations. B. Shannon Diversity Index visualizing differences and similarities in diversity of OTU composition within samples.

**Fig 5:**
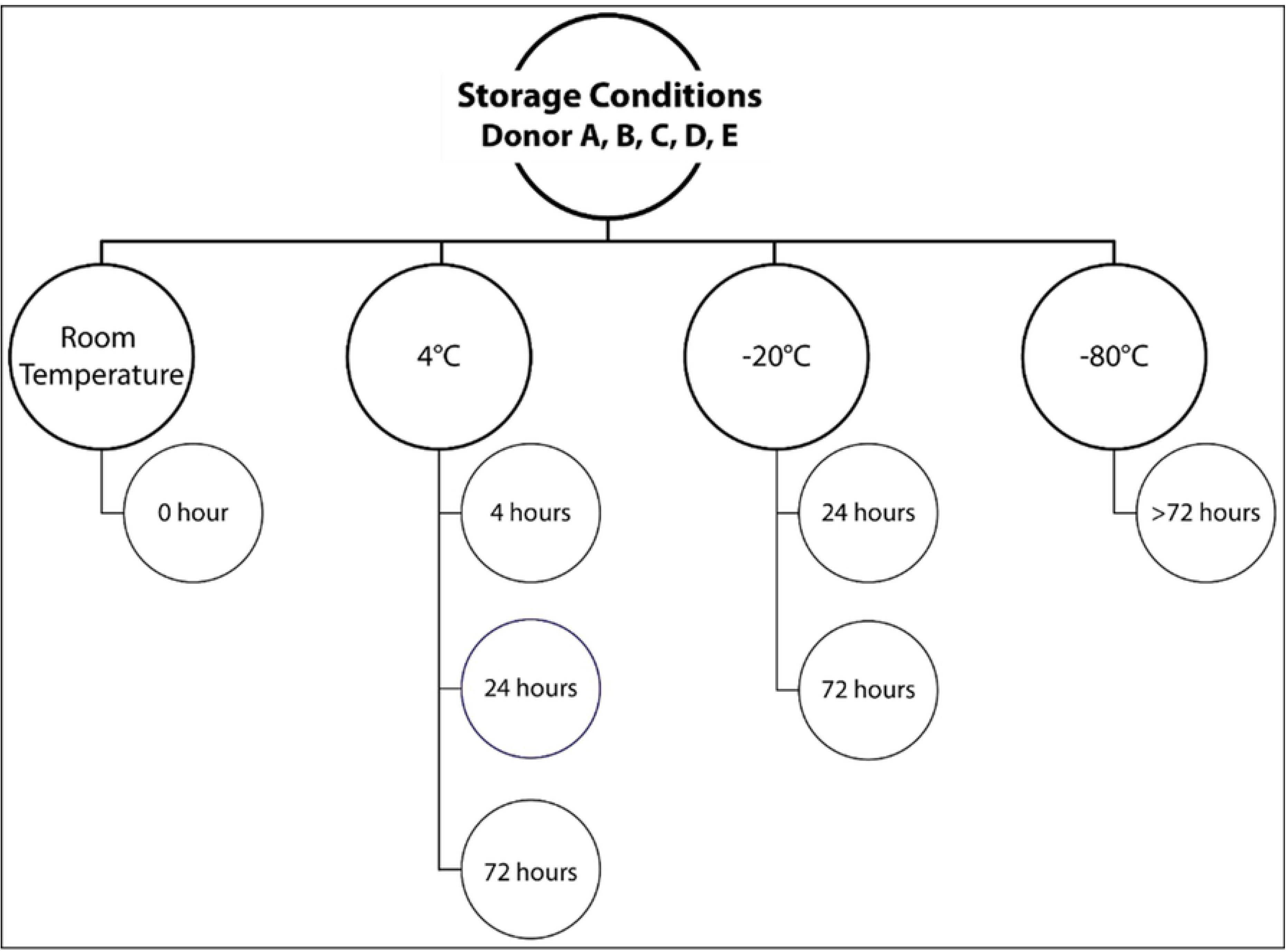
Effects of storage temperature and duration on urinary microbiota beta-diversity.A. Clustering of samples based on PCA of Hellinger transformed OTU abundances. B. Heat map depicting the 20 most common OTUs in different storage conditions. Each name consist of phylum followed by genus name. If no genus could be identified, the best taxonomic assignment is listed.

### Urine microbiota composition is independent of daily and day-to-day variation

First morning urine is often more concentrated than subsequent urine samples throughout the day, while differences in daily routines (e.g. sleep rhythm, diet, sexual activity or exercise) may introduce variations during the day. We therefore speculated that morning urine could contain higher bacterial loads, and possibly a different bacterial composition than urine collected in the evening. To test this hypothesis, we compared urine samples collected in the morning and evening on two independent days from 19 healthy donors (4 women, 5 men, 5 girls, and 5 boys). Following DNA extraction, the resulting DNA yield ranged from <2 to 218.25 ng per mL urine. Importantly, DNA yield did not differ based on within day or day-to-day (data not shown).

Due to the collection method being performed under less controlled conditions (self-sampling by study participants), we expected a higher risk of contamination. In order to avoid false positive samples, we excluded samples that yielded less DNA following the initial PCR amplification for library preparation, compared to the negative controls (0.0538 ng/µL)). 54 of the original 152 samples were below cut-off levels for 1^st^ PCR library amplification. Deselected samples were distributed unevenly as none were from women, 16 (40 %) from girls, 26 (65 %) from men, and 12 (30 %) from boys. The remaining 98 samples (representing 17 participants: 4 women, 4 girls, 4 men and 5 boys) were available for microbiota comparisons. These all showed good sequencing coverage based on a rarefaction curve (data not shown). 16S rRNA gene sequencing resulted in a total of 5,575,050 reads (median 52,232, range 7529 - 122,101 reads per sample) and 2,538 unique OTUs (median 216, range 109 – 469 OTUs per sample) were identified. 93.1 % were assigned to the phylum level and 50.9 % to the genus level.

Mapping of microbiota composition by 16S rRNA gene sequencing did not show any significant difference in OTU richness or Shannon diversity between urine samples collected in the morning or in the evening (Fig 6A and 6B) or on two independent days (Fig 8A and 8B). For beta diversity, we observed that, with the exception of the adult male participants 12 and 15, urine samples maintained similar bacterial compositions regardless of collection time point (Fig 7 and 9). For participant 12, a marked bacterial difference between morning and evening samples was observed, and between morning and evening and weekend and weekdays for participant 15.

**Fig 6:**
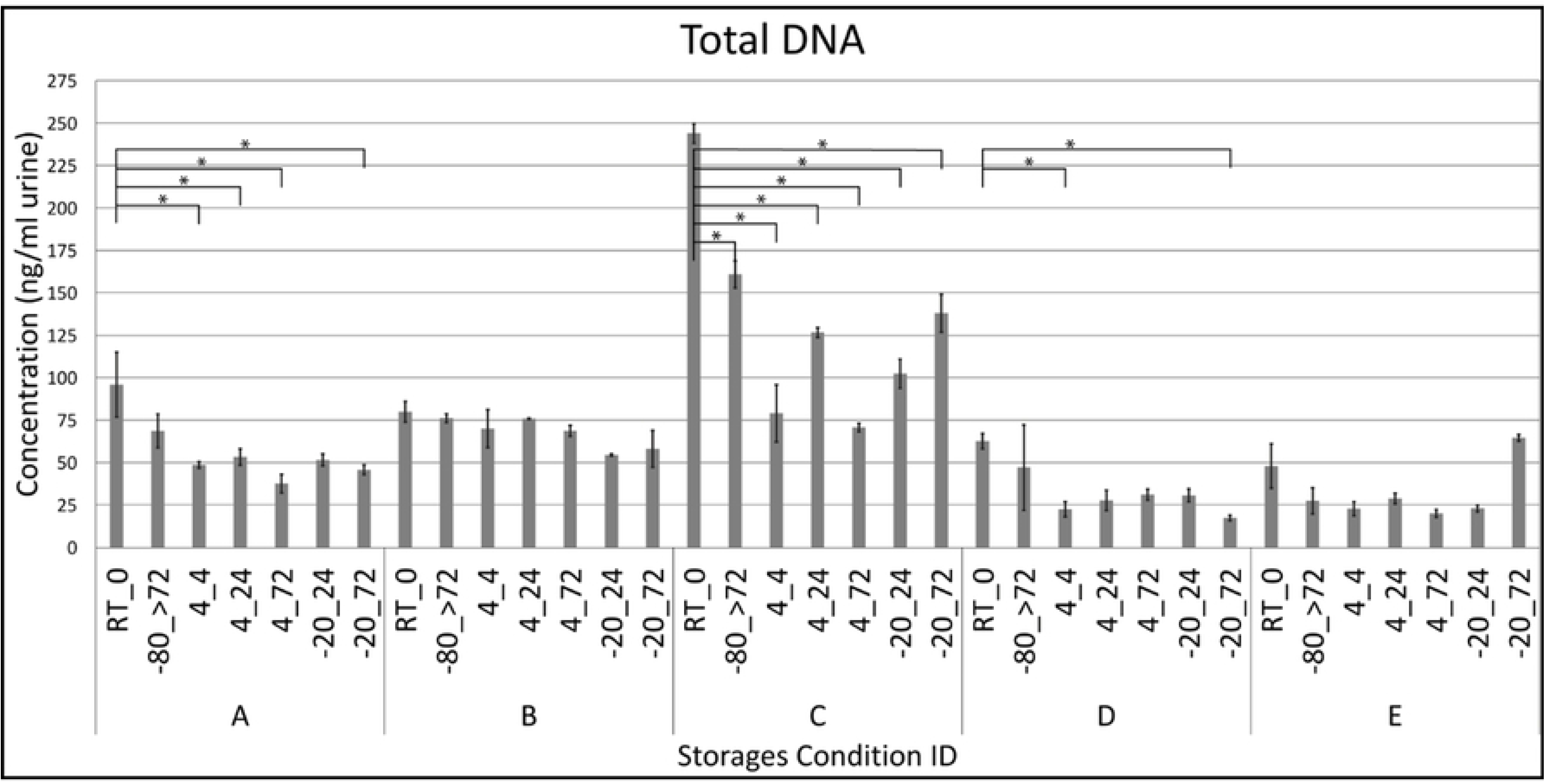
Effects of sampling at evening compared to the following morning on alpha diversity. A. OTU richness depicting number of unique OTUs observed at evening compared to morning. B. Shannon diversity index visualizing similarities in diversity between evening and morning samples. Two donors, one girl (donor 8) and one man (donor 13), were excluded from this part of the analysis, since matching morning-evening sample pairs were not available due to quality control.

**Fig 7:**
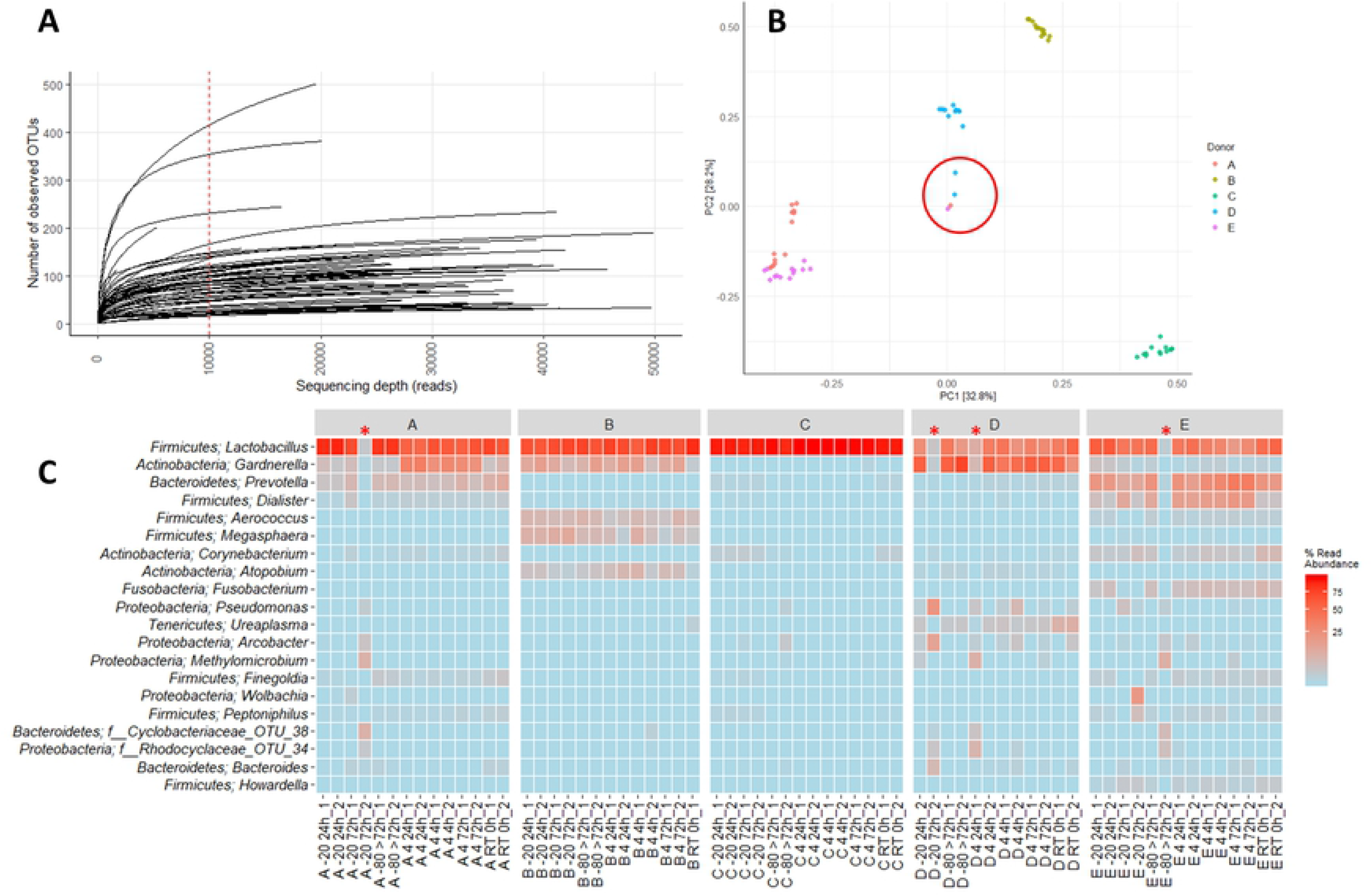
Beta diversity between samples collected in the evening compared to the following morning. A. Relatedness in bacterial composition of samples, as visualized using PCA of Hellinger transformed OTU abundances. Colors indicate donor while shapes indicate time. B. Heat maps listing the 20 most common OTUs for each donor at morning and evening. Each name consists of phylum followed by genus name. If no genus could be identified, the best taxonomic assignment is given. As for figure 7, two donors, one girl (donor 8) and one man (donor 13), were excluded from this part of the analysis, since they did not contain matching morning-evening samples.

**Fig 8:**
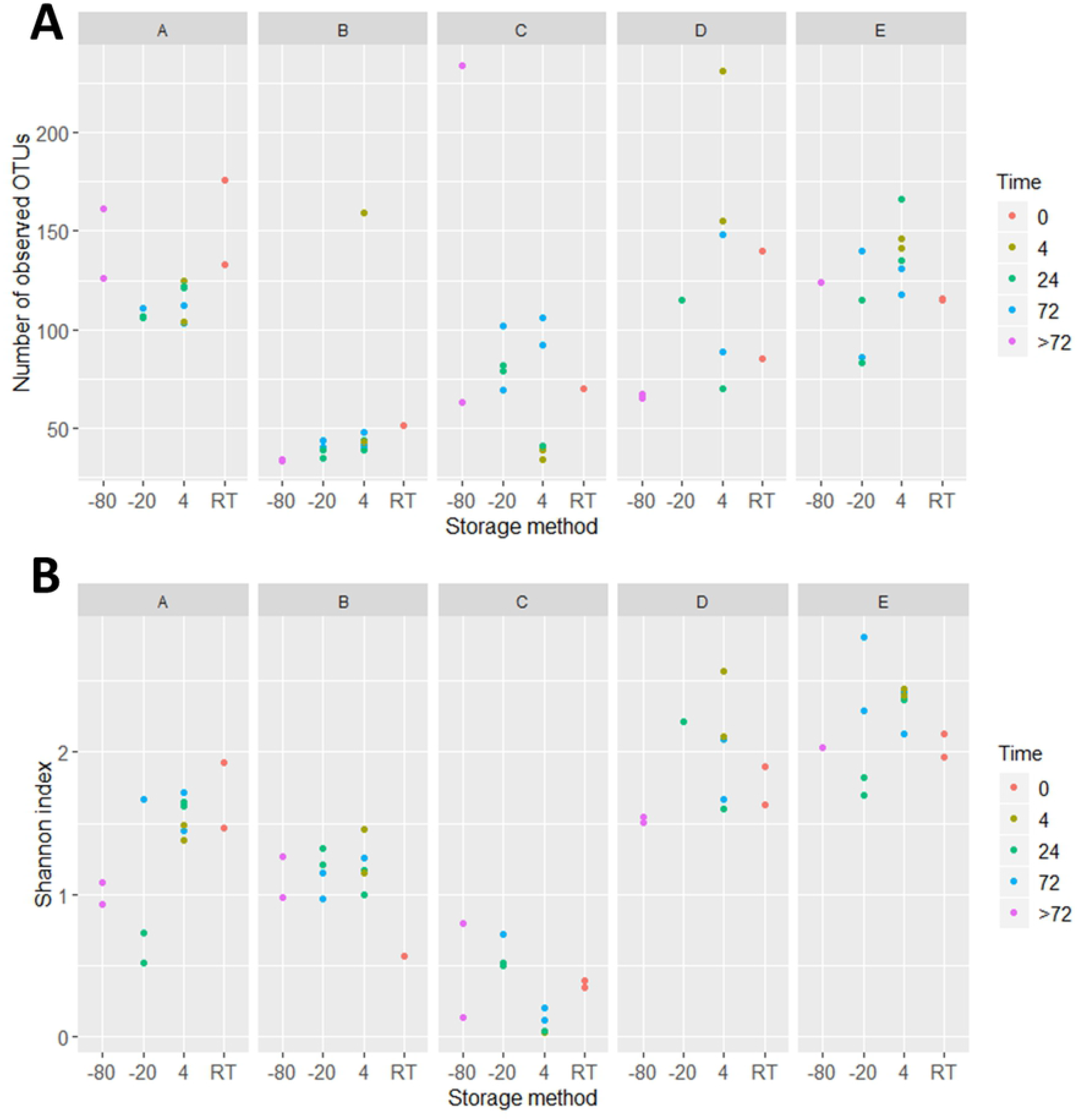
Alpha diversity of urinary microbiota on weekdays compared to weekends. A. OTU richness depicting number of unique OTUs in samples collected during weekdays compared to weekends. B. Shannon diversity index visualizing similarities in diversity of samples collected during weekdays compared to weekends. Three donors, one girl (donor 8) and two men (donors 13 and 15) were removed from this part of the analysis, since they did not contain matching weekday-weekend samples.

**Fig 9:**
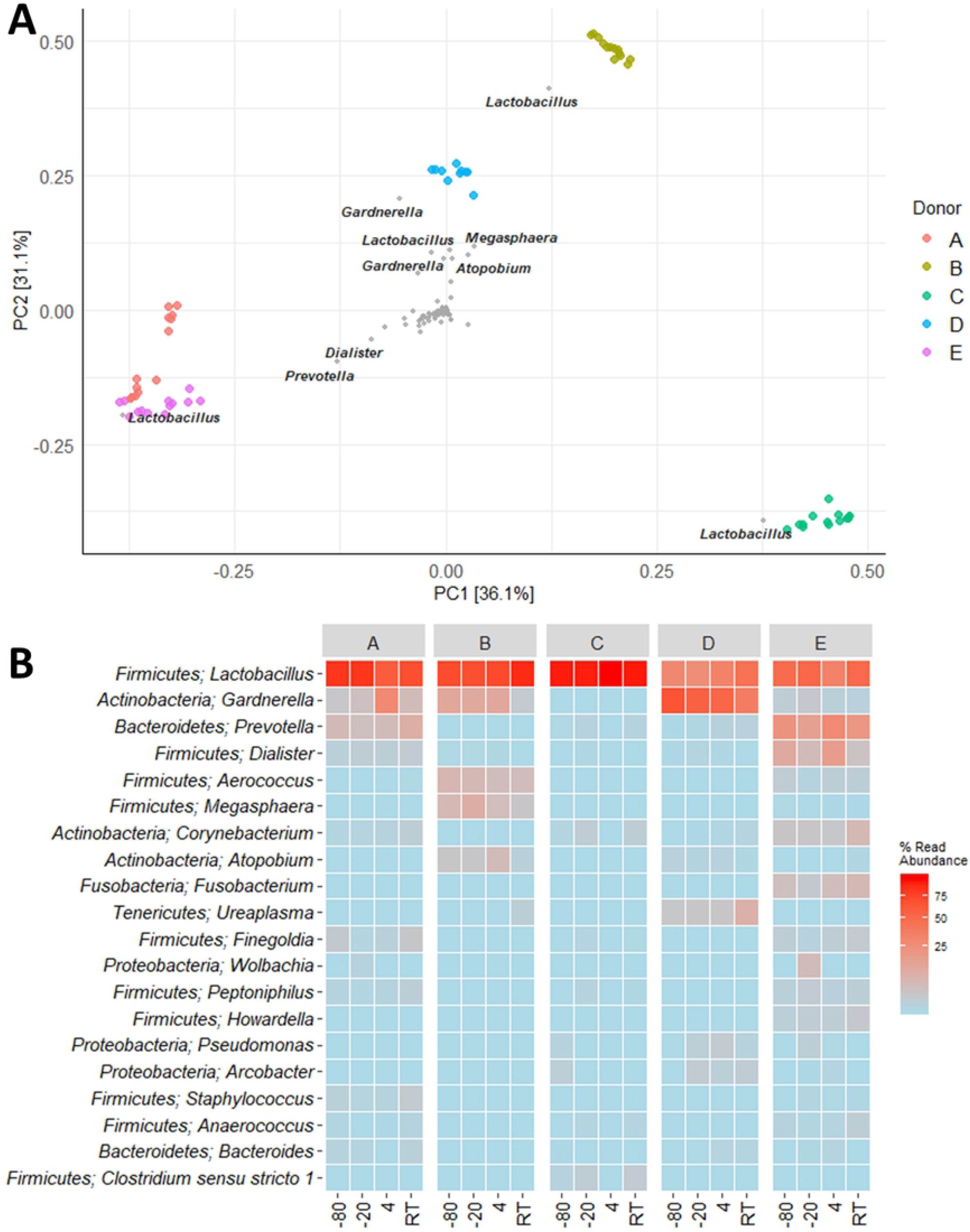
Beta diversity of urinary microbiota at weekdays compared to weekends. A. Variation in bacterial composition of samples collected during weekdays compared to weekends, using PCA of Hellinger transformed OTU abundances. Colors indicate donor while shape indicate day. B. Heat maps listing the 20 most common OTUs for each donor at weekdays compared to weekends. As for figure 9, three donors, one girl (donor 8) and two men (donors 13 and 15) were removed from this part of the analysis, since they did not contain matching weekday-weekend samples.

## Discussion

One of the great challenges, when performing microbiota studies on biological specimens, is always the risk of introducing bias due to methodological vulnerabilities. Studies investigating potential pitfalls are therefore essential to identify confounding factors. Here we show that the urinary microbiota is remarkably stable under different storage conditions and with respect to time.

One major challenge, when investigating the urinary microbiota, is that only low amounts of DNA, and even lower amounts of bacterial DNA, can be extracted from urine[32]. A study by El Bali et al.[32] investigated DNA extraction from urine samples and found that storage temperature had a major impact on DNA output levels. In particular, they showed that storage at −20 °C gave rise to dramatically lower DNA yields compared to samples that were either stored directly at −80 °C or where DNA was immediately extracted. We did not observe the same level of DNA loss in samples stored at −20 °C compared to −80 °C, which may be explained by the longer storage time (15 days) used in El Bali et al. Storage at −20 °C for more than 72 hours may therefore compromise urinary DNA integrity. For gut microbiota studies, maintaining bacterial composition is essential. A previous study by Jung et al.[34] reported, that storage of urine samples without stabilizing buffer at −20 °C or 4 °C, maintained bacterial composition for up to 24 days. This matches the results of our study, allowing collection of urine samples at locations at distance from laboratory facilities.

For samples collected in the home of participants, we observed that a relatively high number of these were excluded due to low DNA yield following the initial PCR during library preparation. Importantly, the samples were not excluded due to lack of high quality sequencing reads, but merely because background contamination could not be ruled out. We have not encountered any urinary microbiota studies that describe exclusion of samples based on background cut-off levels, which leaves the risk of misinterpretation of urinary bacterial loads or profiles.

Importantly, we found that the urinary microbiota composition remained stable across different time points. While microbiota resilience over time has previously been reported in the gut and saliva microbiota[35], this is, to our knowledge, the first study to demonstrate this for the urinary microbiota. This resilience alleviates concerns about timing of urine collection in future studies investigating the urinary microbiota.

Our study suffers from certain limitations. The experiments on the day-to-day and daily variations depended on self-sampling by the study participants in their homes, and are thereby performed in a less controlled environment. This can also be considered as a study strength, since home sampling is an often-used collection method in clinical experiments[36,37]. In fact, we show that the urinary microbiota is stable despite the use of home sampling. Only in two men did we observe inconsistency between morning/evening and weekday/weekend samples. This could be due to biologically relevant fluctuations in microbiota compositions or, more likely, due to contamination from vaginal microbiota of their partner through intercourse prior to urine sampling. In particular, we observe that aberrant microbiota profiles of these men showed similarities to the microbiota profiles of women (e.g. higher relative abundances of *Lactobacillus*). We did not collect data on sexual activity of the participants, which may be considered a potential confounding factor. Another limitation is the large amount of samples that were excluded due to first PCR DNA levels below cut-off values. Whether this fall-out of samples is due to technical issues, e.g. DNA degradation or presence of PCR inhibitors, or due to biological differences in urinary bacterial loads between individuals, is unknown. The latter may be the case since we observed that a very high proportion of samples from men (65 %) did not reach above cut-off levels, indicating that only very little bacterial DNA can be isolated from adult male urine samples. In comparison, none of the samples from women were discarded, 40 % from girls, and 30 % from boys. Our sample size is however too small to make any solid conclusions on age and gender differences in bacterial loads.

The strengths of this study are numerous. We have used a very systematic and structured approach with a relatively large number of participants. Importantly, when evaluating the day-to-day or daily variation, we included study participants of different gender and age. This was to take into account whether there could be differences in time-dependent stability of different core microbiotas. Our data showed that, with the exception of two men, the urinary microbiota was stable regardless of gender and age. Finally, we take into account that very low levels of bacterial contamination may result in false positive samples, leading to misinterpretation of true microbiota profiles.

## Conclusion

In conclusion, we showed that the urinary microbiota is stable over time, and that sub-optimal temperatures for urine storage may be used. We recommend, however, that samples be transferred to −80 °C as quickly as possible after collection, to avoid loss of the already limited DNA in urine samples. In addition, we highly recommend that besides including important clinical parameters such as diagnoses and medication, it is important to consider choice of storage condition and to implement good negative controls and use of background cut-off levels. The latter is especially important when working with low biomass samples. Finally, we recommend studies investigating the effects of sexual activity on the microbiota composition in urine, to determine if this may be a confounding factor. We encourage further studies on the methodological and technical aspects of urinary microbiota research with the aim of providing strong evidence based guidelines.

## Methods

### Study participants and urine collection

In total, 25 healthy volunteers, without symptoms from the bladder (based on self-reporting in a questionnaire prior to study participation) or intake of any antibiotics within the past 3 months, were included into this study. Furthermore, in cases where medicine or hormonal contraceptives were used, these should be taken within the same period on all study days. Use of non-prescription painkillers was not accepted for up to 24 hours prior to urine collection. In addition, the study participants were instructed to avoid urine collection during menstruation, and pregnant women were not included into the study. The identity of all donors was anonymous and no personal data was registered, besides the sex and age interval of which they belonged to. For the initial study on different storage conditions, 5 women were recruited, and for the following study on daily or day-to-day variations, 20 participants were recruited encompassing 5 men (18-50 years), 5 women (18-50 years), 5 boys (5-10 years), and 5 girls (5-10 years). One woman was however excluded from the latter of the two studies, due to incorrect storage of the collected urine sample, leaving 19 participants in total.

For the study on storage conditions, urine was collected at the laboratory by the clean catch method. Samples were immediately aliquoted in tubes with 10 mL urine and transferred to the specified storage conditions as summarized in Fig 1, or subjected directly to DNA extraction (RT sample). All conditions were tested in duplicates (two aliquots from each urine sample). For samples stored at −20 °C, a freezer corresponding to a domestic freezer was used to mimic a home collection situation. All samples were finally collectively stored at −80 °C, to rule out bias due to differences in low temperature exposure, until further processing.

For the study on daily or day-to-day variations, the participants collected urine at home by the clean catch method, and urine samples were immediately transferred to −20 °C domestic freezers. All participants delivered two first morning samples (weekday and weekend) and two evening samples (weekday and weekend) and likewise, collections on a weekday or weekend day were represented by two independent samples (morning and evening). This gave rise to two independent samples per time point from each participant. Each independent sample was furthermore divided into two aliquots for duplicate DNA purification. Children were assisted by a parent to ensure correct sampling. Samples were subsequently, within 24 hours, transported on ice to the laboratory. Upon arrival to the laboratory, the samples were stored at −80 °C until further processing.

### Ethics statement

Oral consent was obtained from all study participants, or from parents or other legal guardians if participants were below 18 years of age. The Regional Ethical Committee of Northern Denmark reviewed the study protocol. Since no personal information were collected from study participants and no intervention was performed, the Ethical Committee judged that no further approval was required.

### DNA purification

Bacterial DNA was isolated from 10 mL of urine using the QIAamp Viral RNA Mini Kit (Qiagen) according to manufacturer’s recommendations. Prior to DNA extraction, urine samples were centrifuged at 3,000x*g* for 20 minutes. Pellets were resuspended in PBS, lysis buffer added, and a bead beating step was included using the TissueLyser LT (Qiagen) for 2 minutes at 30 Hz with a 5 mm stainless steel bead. DNA yield was measured by the NanoDrop™ Lite Spectrophotometer (Thermo Fisher Scientific) or by fluorometric quantification using the Qubit 4 Fluorometer (Thermo Fisher Scientific) together with the Qubit dsDNA HS Assay Kit (Thermo Fisher Scientific).

### 16S rRNA gene sequencing

Bacterial 16S rRNA gene sequencing targeting the V4 region, was performed by DNAsense (Denmark), and followed a modified version of an Illumina protocol[38], as described by Albertsen et al[39], with an initial amplicon PCR. Due to upgrades in primer design during the experiment, different versions of reverse primers targeting the V4 region of the 16S rRNA gene were utilized in the different parts of this study. For the study on storage conditions, the following primer sequences were utilized (Forward: 5’-GTGCCAGCMGCCGCGGTAA-3’, reverse: GGACTACHVGGGTWTCTAAT), while for the study investigating variations between evening and morning and day-to-day, a slightly modified reverse primer was used (5’-GGACTACNVGGGTWTCTAAT-3’). Samples were pooled and sequencing was performed on a MISeq (Illumina, USA), as previously described[38]. To measure error rate during sequencing and batch effects, a 20 % PhiX control library was added. As negative control, nuclease-free water was used, while a complex sample obtained from an anaerobic digester system was utilized as a positive control.

### Bioinformatics and statistics

Quality of sequencing reads was analyzed using FastQC (Babraham Bioinformatics, UK). Forward reads were quality trimmed using Trimmomatic v 0.32[40] to produce reads with a Phred score of at least 20 and a length of 250 bp. Subsequent bioinformatics followed the UPARSE workflow[41] to remove chimeras, cluster OTUs based on 97 % identity and assign taxonomy using the RDP classifier as previously described[26].

Data analysis was performed in R version 3.5.3[42] through the Rstudio IDE (http://www.rstudio.com/) using the ampvis2 package v.2.4.5[39], as well as Microsoft Office Excel 2013. To evaluate sequencing coverage, a rarefaction curve was generated using the amp_rarecurve command, while DNA quantity following the initial PCR amplification was compared to the corresponding negative control to rule out background contamination. Alpha-diversity was determined using OTU richness and Shannon Diversity Index, as implemented in the amp_alphadiv command. Beta-diversity was determined using PCA clustering and heat maps. PCA of variance in Hellinger transformed OTU abundance between samples and storage conditions were determined using the amp_ordinate function, while heat maps displaying the most commonly found OTUs in differing conditions, were produced using the amp_heatmap function. For continuous data like DNA concentration, distribution was tested using Shapiro-Wilks test while variance was tested using Bartlett’s test. Normal distributed data was expressed by mean values and analyzed using ANOVA followed by Bonferroni post-hoc test, while data that was not normal distributed or did not have equal variances, was expressed as median values and analyzed using Kruskal-Wallis Test followed by Dunn’s post hoc test. Differences were considered statistical significant for p<0.05.

## Supplementary information

**S1_Metadata CP154**. 16S rRNA gene sequencing metadata. Information describing the samples used for investigating storage.

**S2_otus CP154**. 16S rRNA gene sequences. FASTA file containing the DNA sequences generated using 16S rRNA gene sequencing of the samples used for investigating storage.

**S3_otutable CP154**. 16S rRNA gene sequencing OTUtable. OTU table documenting numbers of different OTUs generated during the 16S rRNA gene sequencing of the samples used for investigating storage.

**S4_metadata CP303**. 16S rRNA gene sequencing metadata. Information describing the samples used for investigating bacterial variation across different time of day or between weekday or weekend.

**S5_otus CP303**. 16S rRNA gene sequences. FASTA file containing the DNA sequences generated using 16S rRNA gene sequencing of the samples used for investigating bacterial variation across different time of day or between weekday or weekend.

**S6_otutable CP303**. 16S rRNA gene sequencing OTUtable. OTU table documenting numbers of different OTUs generated during the 16S rRNA gene sequencing of the samples used for investigating bacterial variation across different time of day or between weekday or weekend.

